# A Nature-based solution in practice: ecological and economic modelling shows pollinators outperform agrochemicals in oilseed crop production

**DOI:** 10.1101/628123

**Authors:** Rui Catarino, Vincent Bretagnolle, Thomas Perrot, Fabien Vialloux, Sabrina Gaba

## Abstract

Nature-based agriculture, reducing dependency on chemical inputs, requires using ecological principles for sustainable agro-ecosystems, balancing ecology, economics and social justice. There is growing evidence that pollinator-dependent crops with high insect pollination service can give higher yields. However, the interacting effects between insect pollination and agricultural inputs on crop yields and farm economics remain to be established to reconcile food production with biodiversity conservation. We investigated the effects of insect pollination and agricultural inputs on oilseed rape (*Brassica napus* L.). We show that not only yield but also gross margins are 16-40% higher in fields with higher pollinator abundance than in fields with reduced pollinator abundance. This effect is however strongly reduced by pesticides use. Higher yields may be achieved by either increasing agrochemicals (reducing pests) or increasing bee abundance, but crop economic returns was only increased by the latter, because pesticides did not increase yields while their costs reduced gross margins.

## Introduction

Achieving world food production to meet the demands of a growing population while minimizing environmental impacts is a major challenge [1]. Modern agriculture may be at a tipping point, with nature’s supporting mechanisms failing [2] and artificial inputs such as fertilizers and pesticides being either ineffective or used inefficiently [3,4]. There is also growing recognition that ecosystem service degradation is not only an environmental problem but has huge economic consequences [5]. The next key challenge in western agriculture is, therefore, to stabilize crop yields while decreasing the dependence on agrochemical inputs [6]. Nature-based solutions for agriculture are a key EU research target [7] and form the basis of agro-ecology [6]. This requires using ecological principles for sustainable agro-ecosystems, balancing ecology, economics and social justice [8]. Sustainable agro-ecology relies on maximizing the replacement of agro-chemicals by natural capital and ecosystem functions, while minimizing the reduction in yield and increasing farm profitability.

Insect pollination is a key intermediate ecosystem service as a third of human food production benefits directly or indirectly from it [9]. However, in recent years, the abundance and diversity of insect pollinators have been declining worldwide, affecting pollination services [10,11]. At the same time, the cultivated area of oilseed rape (OSR, *Brassica napus* L.) is rapidly increasing, driven by increasing demand, so that OSR production may become limited by pollinator abundance such as honeybees [12]. Pesticides are used in large quantities for intensive farming to mitigate the direct impact of pests or weeds on OSR yield [13–15], but these pesticides, and especially insecticides, can increase the mortality rates of pollinators [16] and reduce their efficiency [17–19]. Herbicide, by modifying weeds abundance in crops may positively [20] or negatively [21] also influence pollinator abundance.

OSR is considered to be both self-pollinated and wind-pollinated [22]. Though, insect pollination can increase the yield of winter OSR by 20-35% [23,24], with a possible benefit of €2.6 M.year^−1^ for the whole of Ireland [25]. Estimating the extent to which OSR production relies on insects for pollination services is, however, less easy than usually thought [26,27]. Firstly, measuring pollination services by quantifying the reduction in yield when pollinators are excluded, also excludes other ecosystem services and may stress the plants [26]. Secondly, the benefits of a pollination service in terms of increased yield is often assumed to be independent of the level of inputs [26,28]. However, crop production is a complex multi-scale system [29,30] which involves inputs that may interact with abiotic factors (e.g. soil properties), the biodiversity, and the services they provide. Recent studies have demonstrated that the value of insect pollination depends on the soil fertility [30,31], field size [32] and farming practices such as the selection of cultivars [22] and pest control [33]. When pollinators are limited, farmers can change their practices to compensate for poor pollination by, for example, increasing fertilizer applications [34]. Thirdly, pollinator abundance and pollination efficiency vary with the composition of the surrounding landscape [35]. Landscapes with large quantity of pollinator-friendly areas, such as semi-natural habitats (SNH: woodlands, meadows) which can increase the abundance of pollinators [28] or attract pollinators away from the OSR fields [36]. Recent research [37] has showed that a higher proportion of OSR in the surrounding landscape may also decrease insect pollination by spatial dilution of the pollinator population. Moreover, pollinator abundance decreases with distance from the edge of an OSR field [38] especially for wild pollinators with limited range [39]. Overall, the extent to which pollinators and other farming practices interact to increase or limit OSR yields remains little known [29,30].

Although OSR is perhaps the most well studied crop regarding the interaction between pollination services and farmers practices, very few studies have been performed under real working farm conditions (but see for exception Lindström studies in Sweden [40,41] and Perrot et al. (2018)[24]). Moreover, studies generally investigated the effect of a single farming practice on the contribution of pollinators, such as fertilizer inputs [30,34], insecticide use [41], pest exclusion [33] or cultivar type [34,40]. Furthermore, the effect of interactions between pollination and farming practices on farm income (Fig. 1) have never been investigated, despite pollination being one of the most commonly assessed services. Existing studies of the economic value of pollination have been almost exclusively illustrative, with few cost-benefit analyses of the role of pollinators (review in Hanley *et al.* [42]). In our study, we address this gap by quantifying the effect of bee visitation on yields and gross margins for OSR with diverse farming practices and landscape characteristics (Fig. 1). We collected the data over six years from 294 OSR fields along landscape gradients with varying proportions of arable and semi-natural habitats (SNH), ensuring a wide variation in pollinator abundance and diversity (the pollinators were counted in the focal fields). We used linear models fitted to this large dataset to quantify the individual and combined effects of farming practices, soil quality and, bee abundance (on a subset of data), on OSR yield and gross margin. We then used the model to test the effect of maximizing pest control or bee abundance on yield and gross margin. We predicted that reducing herbicide (presumably increasing weed abundance) would increase the attractiveness of the OSR field and the bee visitation rate (this assumes no competition for pollinators between higher weed abundance and OSR plants). We also predict that reducing insecticide (presumably decreasing the bee mortality rate) use would not only increase OSR fruiting success and yield, but increase the gross margin further by reducing costs. Our findings provide an important contribution to the evidence-based promotion of biodiversity as a means of increasing yield and farming profit, an essential step for the adoption of nature-based solutions.

**Figure 1:**
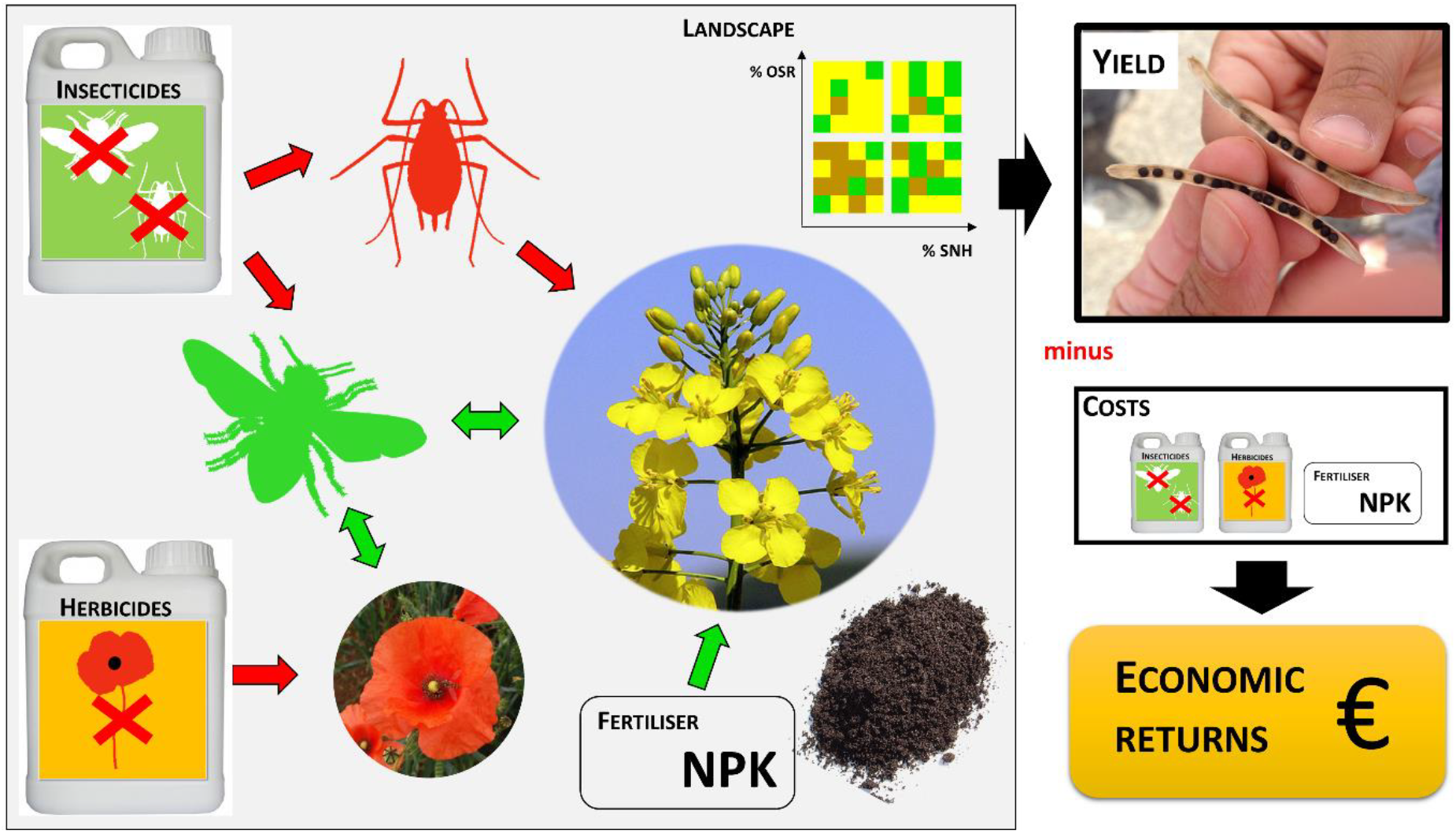
Schematic representation of the relationships between soil type, agricultural practices, bees, landscapes and their effect on yield and economic returns. Red arrows indicate negative interactions, whereas green ones show positive ones.

## Material & methods

### Study area

The study took place from October 2011 to August 2016 in the LTSER “Zone Atelier Plaine & Val de Sèvre”, a long term social-ecological research site covering 450 km^2^ [43] in central western France (46.23°N, 0.41W). It is an agricultural landscape dominated by intensive cereal production, with 8-12% OSR, and average field size of 4-5 ha. The site is also used by professional or amateur beekeepers who own several hundreds of hives, though none of them contract or are paid by farmers for crop pollination. Information about crop yields and farming practices (pesticide and fertilizer use, tillage and mechanical weed control) and general information about the farm (number of crops, agricultural equipment) were collected by farm surveys after harvest. The sample comprised 142 farmers with 294 OSR fields of which 273 fields were sown with hybrid OSR and 21 with pure line OSR (further details on field selection in electronic supplementary material, methods S1). The large majority of farmers (103) managed two fields (2.1±1.4 fields per farmer), and nineteen farmers managed four or more fields. The field size ranged from 0.4 ha to 28.5 ha (mean 6.9±5.0 ha). The soil type varied from very poor dry soil 20 cm deep or less, to 50 cm silt, and was classified in four categories: three highly calcareous soils, with depths of 20, 30 and 40 cm, and one with red silt over limestone.

### Insect pollinator surveys

Between 2013 and 2016, the abundance and diversity of the major groups of flower-visiting insects, including bees (Hymenoptera, Apoidea, Apiformes) and hoverflies (Diptera, Syrphidae) were surveyed [44]. A total of 85 fields (10, 19, 24 and 32 in 2013, 2014, 2015 and 2016) were sampled using both pan traps and sweep nets to get local estimates of the pollinator abundance and richness. The counts of four groups of pollinators (honeybees, bumblebees, other wild bees, and hoverflies) in each field obtained by these, and were combined to provide pollinators abundance index (further details in electronic supplementary material, Methods S2). Due to their limited effect in the study area as demonstrated in [24], hoverflies were excluded from the calculation of pollinator abundance. For each three remaining groups of pollinators and for each field, we averaged the counts for each trapping method. Then, we standardized the values using z-scores [45] across the whole sample size per trapping method. The z-scores for pan-traps corresponded to the total abundance catch per field which were centred (mean of total abundance are removed to each value of total abundance) and reduced (each total abundance value are divided by the standard deviation of total abundance). The final total abundance for each three groups of pollinators was the sum of z-scores for sweep net and pan traps counts in 2013 and 2014, and for visual counts and pan traps in 2015 and 2016. This first metric was called total pollinator abundance. A second metric was further derived, since in our study area, the main bee pollinators in OSR fields are by far *Lasioglossum* spp. (a wild bee) and honeybees [24]. We thus used the sum of the reduced-scores values of these two species/genus as a bee index (electronic supplementary material, Table S1).

### Farm surveys

The general farm statistics obtained from the survey questionnaires during interviews are given in electronic supplementary material, Table S2. From these surveys, we derived the treatment frequency indicator (TFI) as a pesticide use indicator. TFI is a quantitative index that measures the intensity of applications as the number of dose applied per unit of cropped area in relation to the recommended dosage per crop type [46]. TFI reflects the recommended dose necessary to control pests and can be broken down per group of pesticides (herbicide, insecticide and fungicide) or aggregated for all pesticides. TFI per hectare is expressed as:

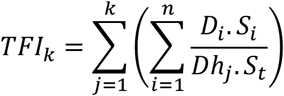

where *D*_*i*_ is the dose in application *i*, *Dh*_*j*_ is the national approved dose for pesticide *j*, and *S*_*i*_ is the surface area treated in application *i* and *St* is the total field area [47]. This includes all the pesticide treatments applied in a given crop field. The recommended dose is defined for each combination of pesticide product and crop type. We computed for each field a global TFI and a TFI for each group of pesticides. A TFI equal to one, e.g. for herbicides, means that the farmer either: (i) applied a single product at the recommended dose in the entire field; (ii) applied two products at half of their recommended dose; or (iii) a single product applied twice at the recommended dose on only half of the surface of the field. For our sample of farms, the global TFI varied from 0.6 to 11.3 (mean: 4.9±1.8, N=294).

Since the inorganic nitrogen in mineral fertilizers is rapidly available to plants, the quantity of nitrogen used was directly calculated from the fertilizer composition and the quantity applied. However, organic compounds with nitrogen are relatively stable and must be mineralized to be available to the crops. The quantity of nitrogen mineralized in organic fertilizers was calculated using the method described by Jeuffroy and Recous [48].

### Statistical analyses

Using the complete dataset (294 fields), we first analysed with a linear mixed model (LMM), the effects of farming practices (fertilizer and pesticides) and soil type (four class) on both yield and gross margin (GM; for further details on gross margins calculation, see electronic supplementary material, Methods S3) accounting for direct and interacting effects. We included interactions between practices (fertilizers) and soil types to account for farmers adapting their practices to soil quality. We also included Farmer ID as a random factor to account for varying number of fields per farmer, and present results in the proportion of variance explained by the fixed factors (marginal R^2^, R^2^m), and the one explained by both the fixed and random factors (conditional R^2^, R^2^c). To estimate the effect of pollinators, we then added bee abundance index and its two-way interactions with the farming practices. The effect of pollinators was studied only for years 2013 to 2016 with a sample size of 85 fields, as bees were not sampled before 2013. In this dataset, since 80% of farmers managed only one field, Farmer ID was not included as a random factor. Finally, we added field size and landscape metrics to the model. The landscape was modelled as the percentages of OSR and SNH (meadows, woodland and hedges, considering a hedge to have a width of two meters), outside the focal field at eight buffer sizes, from 250m to 2000m. Buffer distance was measured from the focal field edge, not the centroid, because the field size was highly variable. The model with buffer width with the highest explanatory power was kept (see below). All models were checked for normality and homoscedasticity. Collinearity was low in all models, with variance inflation factors (VIF) less than 3.1.

At each step, we selected the linear models and linear mixed models with the highest explanatory power, using a multi-model Akaike information criterion method and model averaging using the “dredge” function in MuMin R package [49]. The model averaging approach provides an estimate of the uncertainty of each coefficient [50]. We kept all models with AIC less than 2.0 greater than the best model [50]. The average model was considered to be the best explanatory model. Consequently, although similar set of variables were included in the models for yield and gross margin, after the model selection procedure, different set of variables can be retained. The weight of a parameter was calculated as the sum of the Akaike weights over all of the models in which the parameter was retained [50]. The total amount of variance explained (R^2^) was calculated using the model with the smallest AIC among all models in which the parameter was retained. Farming practices were also standardized per year using z-scores. This transformation does not constrain the variability found in the raw data and allows focusing on each effect independently of the year effect.

Based on our empirical data, we finally explored whether the losses due to reducing the use of herbicides and insecticides could be balanced by an increase in the yield and/or GM due to an increase in bee abundance. We choose to analyse the sum of herbicides, insecticides and fungicides, combining them into a single pesticide TFI, i.e. the sum of each individual TFI. We then used a LM including pesticide TFI, bee abundance and the interaction between bee abundance and pesticide TFI. Annual variation in yield was taken into account by subtracting the average yield of the studied year. We varied the TFI for pesticides and the bee abundance within the observed range of values assuming that the pressure from pests was not increased by the reduction in insecticide. To test the robustness of this assumption, we assessed the relationship between OSR yield, insecticide use and insect pest abundance, using a LM fitted to a third dataset with 74 data points over three years (18 in 2014, 24 in 2015 and 32 in 2016) for which insect pest abundances were available. The effect of pesticides on insect pest abundance was tested using a linear model with insecticides, herbicides and fungicides as explanatory variables. Pest abundance was obtained from the pan trap surveys which give good predictions of pest abundance in OSR inflorescences [51] (see electronic supplementary material, Methods S4 and Table S3).

## Results

### Effect of farming practices on OSR yields and gross margins

Overall, OSR crop yield averaged 3.1 t.ha^−1^ (±0.6, range 1.6:5.4, n=294), red soils showing a significantly higher yield (c. 16% on average) than the other soil types. We tested whether farming practices (fertilizer and pesticide use), soil type and the two-way interactions between fertilizer and soil type affected yield using the complete dataset. The best model (explaining R^2^m = 13.98% of the variance and R^2^c = 46.87%) showed that fungicide significantly increased yield (Table 1a). For gross margin (GM), all inputs were kept in the selected model, as well as all interactions between fertilizer and the soil type (explaining R^2^m=36.37% of the variance (R^2^c=48.72%), Table 1b). The practices most affecting yield and GM were quite different. But most importantly, except phosphorus, all inputs kept in the final model negatively affected GM (Table 1b), including the significant negative effect of nitrogen and herbicides. The soil type and its interaction with nitrogen also had significant effects, with more effect for red soils. Keeping only the variables selected for the yield model (Table 1a) resulted in a model with slightly poorer fit and fewer explanatory variables (ΔAIC=163.17, R^2^m=14.97% and R^2^c =40.96%). Our results further suggested that neither insecticides nor herbicides had a direct significant effect on yield (Figs. 2a, b), but both strongly reduced gross margins (Figs. 2c, d).

**Table 1.** Models of yield (a) and gross margin (b) as a function of farming practices and soil type and their interaction. Weight (w), estimated coefficient (β), 95% confidence intervals (CI) and *p-*value are given for each explanatory variable for the average yield and GM models. β and CI are not given for categorical variables. Significant terms with confidence intervals not including zero are in bold. All explanatory variables were centred/reduced before analysis.

**a) Yield**
| | w | $\beta$ | Lower CI | Upper CI | *p*-val |
| --- | --- | --- | --- | --- | --- |
| **Fungicides** | 0.34 | 0.081 | 0.0143 | 0.1479 | **0.0173** |
| **Soil type** | 1.00 |  |  |  | **<0.0001** |

**b) Gross Margin**
| | w | $\beta$ | Lower CI | Upper CI | *p*-val |
| --- | --- | --- | --- | --- | --- |
| **Nitrogen** | 1.00 | -132.02 | -278.77 | -136.84 | **0.0026** |
| Potassium | 1.00 | -65.75 | -139.36 | 37.10 | 0.2051 |
| Phosphorus | 1.00 | 20.093 | -84.49 | 108.72 | 0.7402 |
| **Herbicides** | 1.00 | -79.61 | -126.27 | -66.69 | **<0.0001** |
| **Insecticides** | 1.00 | -30.52 | -63.85 | -6.97 | **0.0620** |
| Fungicides | 1.00 | -17.86 | -30.93 | 23.03 | 0.2577 |
| **Soil type** | 1.00 |  |  |  | **<0.0001** |
| **Nitrogen x Soil type** | 1.00 |  |  |  | >0.075 |
| Potassium x Soil type | 1.00 |  |  |  | >0.40 |
| Phosphorus x Soil type | 1.00 |  |  |  | >0.35 |

**Figure 2:**
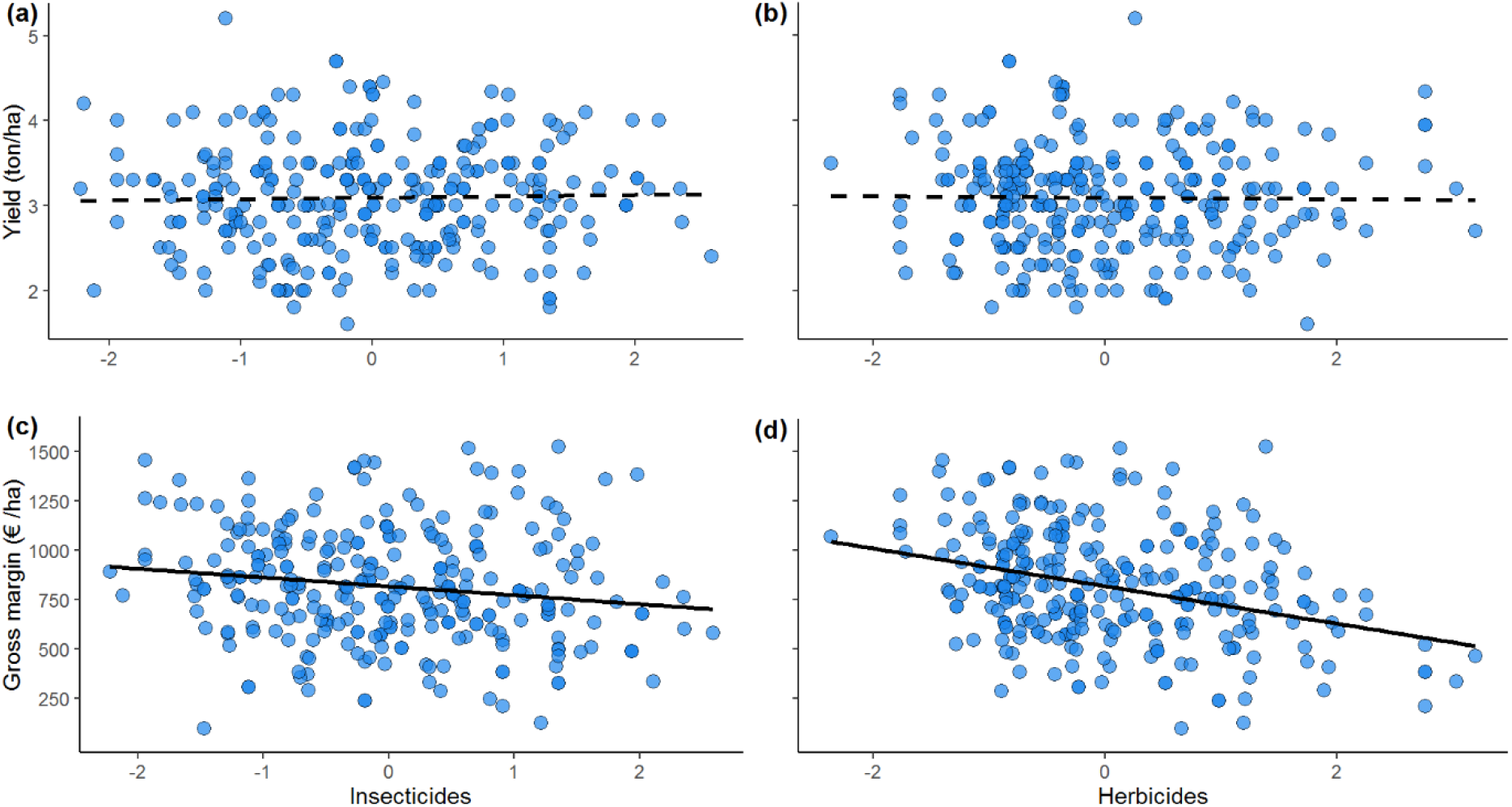
Relationship between insecticides, herbicides on yield (panels a and b) and gross margins (panels c and d), N=294. Solid lines show significant regressions and dashed lines non-significant regressions. Values for both herbicides and insecticides were centred/reduced.

### Effect of bees on yield and gross margin

For yield, adding *Lasioglossum* spp. plus honeybees (i.e., the bee index) improved the model (Table 2). OSR yield increased with bees abundance (*p*-value=0.026, Table 2a), with a significant negative interaction between insecticides and bees (*p*-value=0.039, Table 2a). The model explained 20.6% of the variance (*p*-value<0.01, Table 2a). Including bees removed the soil type effects from the previous model. Although these eliminations might be due to the smaller sample size (85 *vs*. 294), the removal of the soil type was probably due to the higher bee abundance for red soils (about 47% higher, although the difference was not significant, data not shown). Bee abundance and its interaction with insecticide, accounted for about 70.4% of the total variance explained in the yield. Using total pollinator abundance (i.e. including wild bees plus honeybees) did not change the general pattern (electronic supplementary material, Table S4a).

**Table 2.** Models of yield (a), and gross margins (c) as a function of farming practices, soil type, bee index and interactions, and including landscape variables (c). Weight (w), estimated coefficient (β), 95% confidence intervals (CI), and *p-*value are given for each explanatory variable for the averaged yield and GM. β and CI are not given for the categorical variables. Significant terms with confidence intervals not including zero are in bold. All explanatory variables were centred/reduced before analysis. Bees represents the bee index, i.e sum of honeybee and *Lasioglossum* spp. abundances.

**a) Yield**
| | w | $\beta$ | Lower CI | Upper CI | *p*-val |
| --- | --- | --- | --- | --- | --- |
| **Bees** | 1.00 | 0.068 | 0.0081 | 0.1288 | **0.0262** |
| Nitrogen | 0.07 | 0.038 | -0.0824 | 0.1576 | 0.5388 |
| Phosphorus | 0.43 | 0.102 | -0.0322 | 0.2363 | 0.1363 |
| Potassium | 0.24 | -0.070 | -0.1913 | 0.0522 | 0.2625 |
| **Fungicides** | 0.84 | 0.129 | 0.0063 | 0.2525 | **0.0394** |
| Insecticides | 0.55 | 0.047 | -0.0682 | 0.1626 | 0.4231 |
| **Bees x Insecticides** | 0.55 | -0.054 | -0.1035 | -0.0047 | **0.0318** |
| Bees x Fungicides | 0.11 | 0.028 | -0.0215 | 0.0776 | 0.2665 |

**b) Yield, including landscape variables**
| | w | $\beta$ | Lower CI | Upper CI | *p*-val |
| --- | --- | --- | --- | --- | --- |
| **Bees** | 1.00 | 0.077 | -0.020 | 0.215 | **0.0061** |
| Phosphorus | 0.37 | 0.103 | -0.026 | 0.231 | 0.1172 |
| Potassium | 0.12 | -0.073 | -0.070 | 0.161 | 0.2296 |
| **Fungicides** | 0.88 | 0.131 | 0.022 | 0.131 | **0.0305** |
| Insecticides | 0.23 | 0.036 | -0.079 | 0.151 | 0.5406 |
| %OSR | 0.95 | 0.097 | 0.012 | 0.250 | 0.1044 |
| %SNH | 0.17 | 0.045 | -0.193 | 0.462 | 0.4424 |
| **Bees x Insecticides** | 0.23 | -0.047 | -0.093 | -0.954 | **0.0495** |
| Bees x %SNH | 0.05 | -0.038 | -0.089 | 0.014 | 0.1500 |
| Bees x %OSR | 0.63 | -0.052 | -0.112 | 0.009 | 0.0937 |

**c) Gross margin**
| | w | $\beta$ | Lower CI | Upper CI | *p*-val |
| --- | --- | --- | --- | --- | --- |
| **Bees** | 1.00 | 23.95 | 1.319 | 46.582 | **0.0381** |
| Nitrogen | 0.16 | -20.60 | -35.714 | 4.619 | 0.4524 |
| **Potassium** | 1.00 | -69.80 | -119.943 | -19.647 | **0.0064** |
| **Herbicides** | 1.00 | -107.58 | -157.332 | -57.836 | **<0.0001** |
| Insecticides | 0.25 | -20.43 | -12.615 | 39.125 | 0.4310 |
| Bees x Herbicides | 0.46 | -16.27 | -37.733 | 5.200 | 0.1375 |
| Bees x Insecticides | 0.09 | -15.55 | -74.317 | 33.124 | 0.1308 |
| Bees x Potassium | 0.20 | 13.25 | -71.288 | 30.419 | 0.3153 |

The larger field sizes and the presence of other OSR fields nearby may either attract bees or dilute the honeybee population, while *Lasioglossum* spp. may depend on nearby SNH. We thus tested whether including the field size, %OSR and %SNH in the surrounding landscape improved the model. The model that best fitted the data (R^2^ = 22.1%, Table 2b) had a 250 m buffer width. Within this buffer, %OSR and %SNH had a positive effect on yield, although non-significant. All other buffers resulted in lower AIC (data not shown).

For the GM, bee abundance was the only variable with a positive effect (*p*-value=0.0381, Table 2c, Fig. 3a). Farming practices (potassium and herbicide) had a significantly negative effect, and also interacted with bees (Table 2c). Including %OSR and %SNH in the surrounding landscape did not change the effect of pollinators, although the %SNH had a direct significant positive effect for a 250m buffer (electronic supplementary material, Table S5).

**Figure 3:**
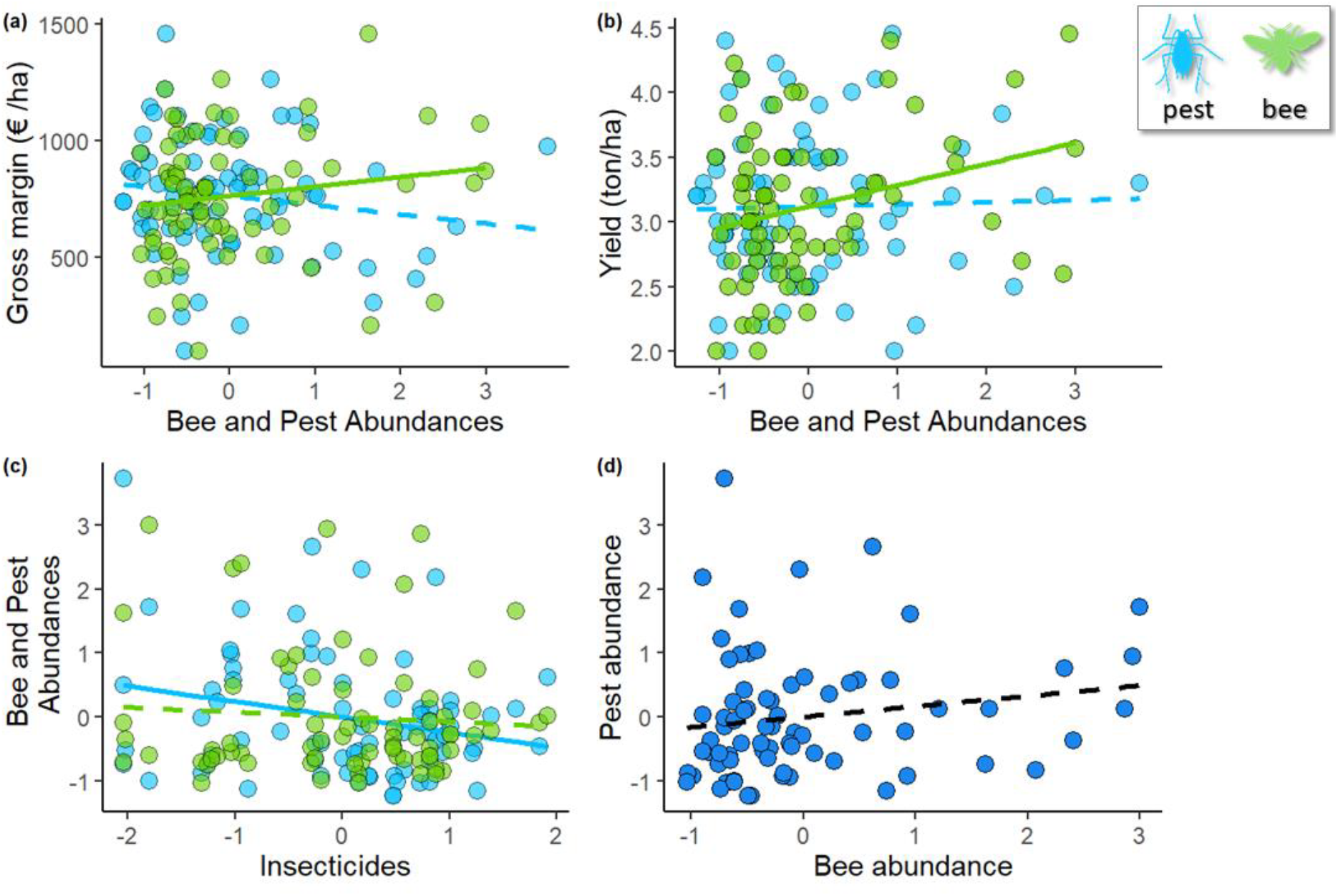
Plot a) shows the effect of pest (blue dots) and bee (green dots) abundance on gross margins; plot b) shows the effect on yield; plot c) shows the effect of insecticides on bees and pests; plot d) shows the relation between pest and bee abundances. Abundances were centered/reduced. Solid lines show significant regressions and dashed lines non-significant regressions. Bee abundance includes honeybee plus Lasioglossum spp

For average levels of inputs, yield was 0.31 t.ha^−1^ higher and gross margin was 119 €.ha^−1^ (i.e. 16%) higher in fields with the high than fields with the low bee abundance using 0.1-0.9 quantile (Figs. 3a, c). Keeping extreme values of bee abundance (i.e., the lowest compared to the highest) yielded a much larger increase of OSR yield (Fig. 3b; 0.77 t.ha^−1^) and GM (Fig. 3a; 289 €.ha^−1^).

### Trade-offs between pollinators, pesticides and pests to improve gross margins

Since bee abundance had a consistently positive effect on yield and GM, and there was a negative interaction between bee abundance and the use of pesticides, we explored whether higher yields and GM could be obtained by reducing the use of agro-chemicals to increase bee abundance and their contribution to yield. All variables kept in the yield and GM models, except bee abundance, insecticides, herbicides and fungicides were set to their mean values (electronic supplementary material, Table S2). The interactions were visualized using 3D plots with the sum of herbicide, fungicide and insecticide TFIs (hereafter TFI pesticide) on the x-axis, bee abundance on the y-axis and yield or GM on the z-axis. This revealed antagonism between pesticide use and bee abundance, with the latter having a greater positive effect when the use of pesticides was low (Fig. 4). Assuming that the pest pressure remains constant, this antagonism between pesticide use and bee abundance shows that farmers could maximize yield through two opposite strategies: maximizing either pesticide use or bee abundance (Fig. 4a). These strategies, however, had a different effect on GM which was always higher when bee abundance was maximized (Fig. 4b). Additionally, although the use of insecticides reduced the abundance of insect pests (F_1,70_= 5.40, *p*-value=0.023, Fig. 3c), a higher abundance of pests would not significantly affect yield (F_1,70_= 0.08, *p*-value=0.78, Fig. 3b). On the other hand, higher abundance of bees had a strong positive effect on both yield (Fig. 3b) and GM (Fig. 3a). As bees and pests were positively related (though not significantly: *r*_s_= 0.23, *p*-value= 0.23; Fig. 3d), the increase in yield due to the higher bee abundance when insecticide use is reduced, was greater than loss of yield due to the increased abundance of pests.

**Figure 4:**
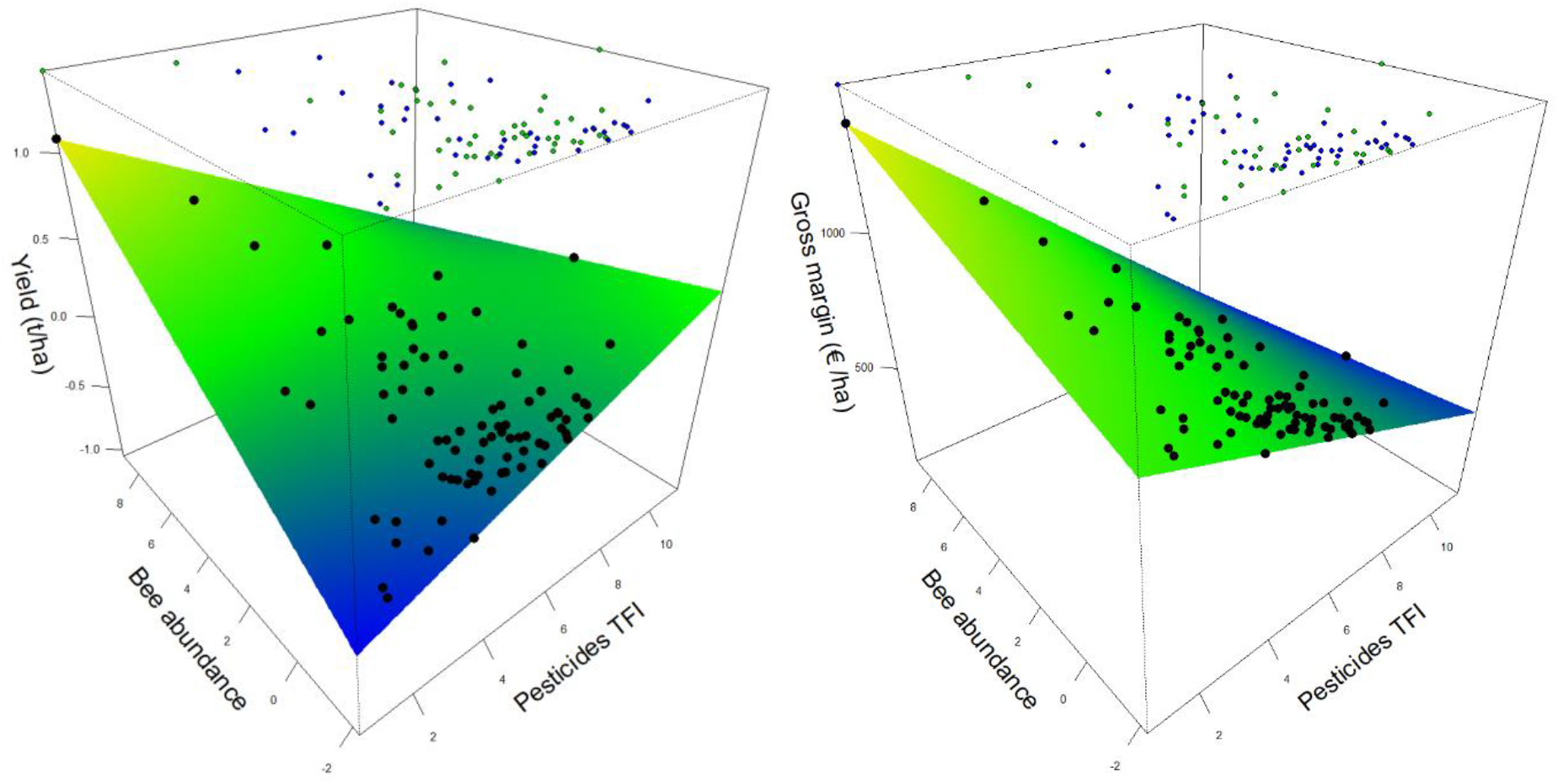
Effect of interaction between bee abundance and the combined herbicide and insecticide TFI on yield (a) and gross margins (b). The green surface shows regions where the yield or gross margin is higher and blue where it is lower. Coloured points represent the raw data points and the black ones predicted values from the model. Positive and negative differences between raw data and predicted values are indicated in blue and green. Bee abundance includes honeybee and Lasioglossum spp, and pesticide TFI is the sum of insecticide and herbicide TFIs. Both explanatory variables were centered/reduced before analysis.

## Discussion

Although ecological intensification appears to be a promising alternative to conventional agriculture (e.g. Pywell *et al.* [52]), there is no consensus on whether it is possible to replace agrochemicals by natural capital and ecological functions without major reductions in yields [53,54]. Insect pollination has been shown to increase OSR yields both in experimental [22,34] and on-farm studies [24,40], but the effect of interactions between pollinators and agricultural practices on yield and income remain largely unknown. Though, the practical implications for farmers, as decision-makers, and for policy-makers are critical [55]. Based on a very large dataset spanning four and six years, this study provides a comprehensive analysis of synergy and antagonism between farming practices and biodiversity, and their effects on yield and income.

Although farming practices overall accounted for about 24% of the variance of the yield, few practices showed significant positive effects. Phosphorous [56] and fungicides [57] were the only inputs with a significant positive effect on OSR yield. Phosphorus may increase OSR yield by increasing the number of pods per plant and seeds per pod [58]. Simultaneously insect pollination, was as well strong determinant of OSR yield, supporting previous experimental studies [23,24,34]. Taking into account farming practices, pollinator abundance explained 50% of the variance of the yield, increasing yields by 0.77 t.ha^−1^ from the lowest abundance to the highest. This is consistent with previous studies that found increases in yield from 0.4 to 1.0 t.ha^−1^ [24,38]. Fertilizer, especially nitrogen, is a recognized driver of yield, but we failed to detect any direct effect of nitrogen fertilizer on OSR yield. Although surprising, the absence of an increase of yield with nitrogen input has already been reported [59,60], and other studies have even reported negative effects [61,62]. This is possibly explained by the ability of modern cultivars to achieve higher yields with lower nitrogen inputs [59]; indeed 93% of the farmers in our study used modern hybrid seed varieties. Our results suggest that, for the farms studied, OSR yield is limited by pollinators rather than nutrient availability [30].

Agricultural practices had little effect on yield which meant that the GM was significantly reduced by nitrogen fertilizer and herbicide applications, as their costs were not recovered in the form of higher yields. Bee abundance was positively correlated with yield, and GM was 15-40% higher with the highest abundance compared with the lowest. This increase of GM assumes that no cost were associated, especially with the presence of hives in the landscapes (i.e. honeybees were dominant pollinator here). In some region, hive rental costs are supported by the farmers. For instance, apple pollination fees are about €40 per hive [63]. Assuming the similar fees per hive for OSR pollination, GM would still be 4%-25% higher with two hives/ha. Very few experimental OSR studies have assessed the economic benefits of pollinators at the field level [26]. Accounting for average production costs per ha, Stanley *et al.* [25] estimated the effect of pollinators on yield in four experimental fields, and then extrapolated to the whole of Ireland to achieve an estimated benefit of €2.6 M.year^−1^. Bommarco *et al.* [23], in a pollination exclusion experiment in ten fields along a landscape gradient, found a 20% increase in the market value of OSR. Our study is the first to assess the financial benefits from pollinators in real farming conditions over 85 fields located along a gradient of pollinator abundance.

The benefits of ecosystem services for crop yield may be affected by agricultural practices such as agrochemical inputs [30,64]. In our study, we focused on the interactions between bee pollination and pesticides. These agrochemicals increase crop yield through decreased insect pests, fungi and weed pressure. However, they can also reduce the benefits of pollination by reducing bee abundance or efficiency, and decreasing the reserves of flowers. With constant insect pest pressure, our analysis showed that higher yields may be achieved by two opposite strategies: increasing agrochemicals (reducing pests) or increasing bee abundance (increasing fruiting success, [24]). But GM was only increased by increasing bee abundance, because insecticides reduced bee abundance and neither insecticides nor herbicides increased yields while their costs reduced gross margins. This result contradicts the dominant arguments about trade-offs between food production and conservation of biodiversity ([65], but see Pywell *et al.* [52]) and shows that nature-based solution can yield to a win-win strategy.

There are two caveats that may limit this interpretation. Firstly, our model assumed constant insect pest and weed pressure, that is, reducing pesticides would not increase their abundances, whereas a reduction in yield may be expected when reducing pesticides [66]. We indeed found that insect pest abundance was lower in fields with high insecticide inputs than in those with low inputs. However, higher insect pest abundances did not translate into reduced yields as there was no relationship between insect pest abundance and OSR yield. It is possible that pest abundance is very low in our study region. For example, with similar trapping method and effort, more of 20 pests were caught in Germany or Estonia [67,68] while only six were caught in our site. It is also possible that OSR plants are able to overcompensate pest damage [69]. However, several recent studies in France have shown that reducing, to a certain extent, pesticides may not reduce yields, as found for herbicides in wheat [4] or for pesticides in general in arable crops [70]. Moreover, pollinators abundance strongly differs between study sites for a same crop type [71], and actually our study region has a particularly rich wild bee community, with more than 250 species [72]. The benefits thus depend on the local pollinator population, part of the natural capital. Further research on the effects of variations in pollinators and farming practices on yields and profits is therefore needed in other agricultural conditions.

New agricultural strategies must be developed to achieve sustainable crop production and reduce dependency on chemical inputs. This study provides a clear demonstration that agro-ecology, by promoting nature-based solutions for agricultural production can be an alternative to conventional agriculture for both food production and farm income. Based on a large-scale field survey, our results therefore support a “win-win-win” balance between crop production, farm income and the environment. The next challenge will be to assess non-market benefits from pollinators to define the value of this natural capital within a landscape, essential for policy-making and land-use planning.

## Authorship

VB and SG designed the study. FV carried out the farm survey. RC and SG performed the analysis, with help from TP and VB. RC and TP wrote the first draft of the manuscript, and VB and SG contributed substantially to revisions and editing. All authors gave their final approval for publication.

## Data accessibility

Upon acceptance of the manuscript, we agree to archive the data in an appropriate public repository.

## Acknowledgements

We would like to express our thanks to Marilyne Roncoroni, Jean-Luc Gautier, Alexis Saintilan and Anthony Stoquert for their help with pollinator trapping and identification. We sincerely thank the farmers of the LTSER “Zone Atelier Plaine & Val de Sèvre” for their involvement on our research programs.

## References

1. Godfray HCJ, Garnett T. 2014 Food security and sustainable intensification. Philos. Trans. R. Soc. B Biol. Sci. 369.

2. Barnosky AD et al. 2012 Approaching a state shift in Earth’s biosphere. Nature 486, 52–58. (doi:10.1038/nature11018)

3. Vitousek PM et al. 2009 Nutrient Imbalances in Agricultural Development. Science (80-.). 324, 1519 LP-1520.

4. Gaba S, Gabriel E, Chadœuf J, Bonneu F, Bretagnolle V. 2016 Herbicides do not ensure for higher wheat yield, but eliminate rare plant species. Sci. Rep., 1–10. (doi:10.1038/srep30112)

5. Sandhu H, Waterhouse B, Boyer S, Wratten S. 2016 Scarcity of ecosystem services: an experimental manipulation of declining pollination rates and its economic consequences for agriculture. PeerJ 4, e2099. (doi:10.7717/peerj.2099)

6. Tittonell P. 2014 Ecological intensification of agriculture — sustainable by nature. Curr. Opin. Environ. Sustain. 8, 53–61. (doi:http://doi.org/10.1016/j.cosust.2014.08.006)

7. EC. 2015 Towards an EU Research and Innovation policy agenda for nature-based solutions & re-naturing cities. Final Rep. Horiz. 2020 Expert Gr. ‘Nature-Based Solut. Re-Naturing Cities’. (doi:10.2777/765301)

8. Altieri MA. 1983 Agroecology: the scientific basis of alternative agriculture. Altieri, M. CRC Press.

9. Klein A-M, Vaissière BE, Cane JH, Steffan-Dewenter I, Cunningham SA, Kremen C, Tscharntke T. 2007 Importance of pollinators in changing landscapes for world crops. Proc. Biol. Sci. 274, 66, 95–96, 191. (doi:10.1098/rspb.2006.3721)

10. Potts SG et al. 2016 Safeguarding pollinators and their values to human well-being. Nature 540, 220–229. (doi:10.1038/nature20588)

11. Garibaldi LA, Aizen MA, Cunningham S, Klein AM. 2009 Pollinator shortage and global crop yield. Commun. Integr. Biol. 2, 37–39. (doi:10.4161/cib.2.1.7425)

12. Breeze TD et al. 2014 Agricultural policies exacerbate honeybee pollination service supply-demand mismatches across Europe. PLoS One 9. (doi:10.1371/journal.pone.0082996)

13. Zhang H, Breeze T, Bailey A, Garthwaite D, Harrington R, Potts SG. 2017 Arthropod pest control for UK oilseed rape - Comparing insecticide efficacies, side effects and alternatives. PLoS One 12, 1–22. (doi:10.1371/journal.pone.0169475)

14. Bijanzadeh E, Naderi R, Behpoori A. 2010 Interrelationships between oilseed rape yield and weeds population under herbicides application. Aust. J. Crop Sci. 4, 155–162.

15. Wang L, Liu Q, Dong X, Liu Y, Lu J. 2019 Herbicide and nitrogen rate effects on weed suppression, N uptake, use efficiency and yield in winter oilseed rape (Brassica napus L.). Glob. Ecol. Conserv. 17, e00529. (doi:https://doi.org/10.1016/j.gecco.2019.e00529)

16. Henry M, Béguin M, Requier F, Rollin O, Odoux J, Aupinel P, Aptel J, Tchamitchian S, Decourtye A. 2012 A common pesticide devreases foraging success and survival in Honey Bees. Science (80-.). 336, 348–350. (doi:10.1126/science.1215039)

17. Stanley DA, Garratt MPD, Wickens JB, Wickens VJ, Potts SG, Raine NE. 2015 Neonicotinoid pesticide exposure impairs crop pollination services provided by bumblebees. Nature 528, 548–50. (doi:10.1038/nature16167)

18. Johnson RM, Dahlgren L, Siegfried BD, Ellis MD. 2013 Acaricide, Fungicide and Drug Interactions in Honey Bees (Apis mellifera). PLoS One 8, e54092.

19. Liao L-H, Wu W-Y, Berenbaum MR. 2017 Behavioral responses of honey bees (Apis mellifera) to natural and synthetic xenobiotics in food. Sci. Rep. 7, 15924. (doi:10.1038/s41598-017-15066-5)

20. Norfolk O, Eichhorn MP, Gilbert F. 2016 Flowering ground vegetation benefits wild pollinators and fruit set of almond within arid smallholder orchards. Insect Conserv. Divers. 9, 236–243. (doi:10.1111/icad.12162)

21. Lander TA, Bebber DP, Choy CTL, Harris SA, Boshier DH. 2011 The Circe Principle Explains How Resource-Rich Land Can Waylay Pollinators in Fragmented Landscapes. Curr. Biol. 21, 1302–1307. (doi:https://doi.org/10.1016/j.cub.2011.06.045)

22. Hudewenz A, Pufal G, Bögeholz A-L, Klein A-M. 2014 Cross-pollination benefits differ among oilseed rape varieties. J. Agric. Sci. 152, 770–778. (doi:10.1017/S0021859613000440)

23. Bommarco R, Marini L, Vaissière BE. 2012 Insect pollination enhances seed yield, quality, and market value in oilseed rape. Oecologia 169, 1025–1032. (doi:10.1007/s00442-012-2271-6)

24. Perrot T, Gaba S, Roncoroni M, Gautier J-L, Bretagnolle V. 2018 Bees increase oilseed rape yield under real field conditions. Agric. Ecosyst. Environ. 266, 39–48. (doi:https://doi.org/10.1016/j.agee.2018.07.020)

25. Stanley DA, Gunning D, Stout JC. 2013 Pollinators and pollination of oilseed rape crops (Brassica napus L.) in Ireland: Ecological and economic incentives for pollinator conservation. J. Insect Conserv. 17, 1181–1189. (doi:10.1007/s10841-013-9599-z)

26. Breeze TD, Gallai N, Garibaldi LA, Li XS. 2016 Economic Measures of Pollination Services: Shortcomings and Future Directions. Trends Ecol. Evol. 31, 927–939. (doi:10.1016/j.tree.2016.09.002)

27. Ouvrard P, Jacquemart A-L. 2018 Agri-environment schemes targeting farmland bird populations also provide food for pollinating insects. Agric. For. Entomol. 20, 558–574. (doi:10.1111/afe.12289)

28. Bartomeus I et al. 2014 Contribution of insect pollinators to crop yield and quality varies with agricultural intensification. PeerJ 2, e328. (doi:10.7717/peerj.328)

29. Seppelt R, Dormann CF, Eppink F V., Lautenbach S, Schmidt S. 2011 A quantitative review of ecosystem service studies: Approaches, shortcomings and the road ahead. J. Appl. Ecol. 48, 630–636. (doi:10.1111/j.1365-2664.2010.01952.x)

30. Garratt MPD, Bishop J, Degani E, Potts SG, Shaw RF, Shi A, Roy S. 2018 Insect pollination as an agronomic input: Strategies for oilseed rape production. J. Appl. Ecol. 0. (doi:10.1111/1365-2664.13153)

31. Tamburini G, Berti A, Morari F, Marini L. 2016 Degradation of soil fertility can cancel pollination benefits in sunflower. Oecologia 180, 581–587. (doi:10.1007/s00442-015-3493-1)

32. Isaacs R, Kirk AK. 2010 Pollination services provided to small and large highbush blueberry fields by wild and managed bees. J. Appl. Ecol. 47, 841–849. (doi:10.1111/j.1365-2664.2010.01823.x)

33. Sutter L, Albrecht M. 2016 Synergistic interactions of ecosystem services: florivorous pest control boosts crop yield increase through insect pollination. Proc. R. Soc. B 283, 20152529. (doi:10.1098/rspb.2015.2529)

34. Marini L, Tamburini G, Petrucco-Toffolo E, Lindström SAM, Zanetti F, Mosca G, Bommarco R. 2015 Crop management modifies the benefits of insect pollination in oilseed rape. Agric. Ecosyst. Environ. 207, 61–66. (doi:https://doi.org/10.1016/j.agee.2015.03.027)

35. Hass AL et al. 2018 Landscape configurational heterogeneity by small-scale agriculture, not crop diversity, maintains pollinators and plant reproduction in western Europe. Proc. R. Soc. B Biol. Sci. 285.

36. Zou Y et al. 2017 Landscape effects on pollinator communities and pollination services in small-holder agroecosystems. Agric. Ecosyst. Environ. 246, 109–116. (doi:10.1016/j.agee.2017.05.035)

37. Holzschuh A et al. 2016 Mass-flowering crops dilute pollinator abundance in agricultural landscapes across Europe. Ecol. Lett. 19, 1228–1236. (doi:10.1111/ele.12657)

38. Woodcock BA, Bullock JM, McCracken M, Chapman RE, Ball SL, Edwards ME, Nowakowski M, Pywell RF. 2016 Spill-over of pest control and pollination services into arable crops. Agric. Ecosyst. Environ. 231, 15–23. (doi:10.1016/j.agee.2016.06.023)

39. Zurbuchen A, Landert L, Klaiber J, Müller A, Hein S, Dorn S. 2010 Maximum foraging ranges in solitary bees: only few individuals have the capability to cover long foraging distances. Biol. Conserv. 143, 669–676. (doi:10.1016/j.biocon.2009.12.003)

40. Lindström SAM, Herbertsson L, Rundlöf M, Smith HG, Bommarco R. 2016 Large-scale pollination experiment demonstrates the importance of insect pollination in winter oilseed rape. Oecologia 180, 759–769. (doi:10.1007/s00442-015-3517-x)

41. Lindström SAM, Klatt BK, Smith HG, Bommarco R. 2018 Crop management affects pollinator attractiveness and visitation in oilseed rape. Basic Appl. Ecol. 26, 82–88. (doi:https://doi.org/10.1016/j.baae.2017.09.005)

42. Hanley N, Breeze TD, Ellis C, Goulson D. 2015 Measuring the economic value of pollination services: Principles, evidence and knowledge gaps. Ecosyst. Serv. 14, 124–132. (doi:10.1016/j.ecoser.2014.09.013)

43. Bretagnolle V et al. 2018 Description of long-term monitoring of farmland biodiversity in a LTSER. Data Br. 19, 1310–1313. (doi:https://doi.org/10.1016/j.dib.2018.05.028)

44. Bretagnolle V et al. 2018 Towards sustainable and multifunctional agriculture in farmland landscapes: Lessons from the integrative approach of a French LTSER platform. Sci. Total Environ. 627, 822–834. (doi:https://doi.org/10.1016/j.scitotenv.2018.01.142)

45. Witte RS, Witte JS. 2010 Statistics. ninth. J. Wiley & Sons.

46. Coll M, Wajnberg E. 2017 Environmental Pest Management: Challenges for Agronomists, Ecologists, Economists and Policymakers. John Wiley & Sons.

47. Kudsk P, Jensen JE. 2014 Experiences with Implementation and Adoption of Integrated Pest Management in Denmark. In Integrated Pest Management: Experiences with Implementation, Global Overview, Vol.4 (eds R Peshin, D Pimentel), pp. 467–485. Dordrecht: Springer Netherlands. (doi:10.1007/978-94-007-7802-3_19)

48. Jeuffroy M, Recous S. 1999 Azodyn: a simple model simulating the date of nitrogen deficiency for decision support in wheat fertilization. Eur. J. Agron. 10, 129–144.

49. Barton K. 2018 Package ‘MuMIn’. CRAN. R-projec., 1–73. See ftp://155.232.191.229/cran/web/packages/MuMIn/MuMIn.pdf (accessed on 2 February 2018).

50. Johnson JB, Omland KS. 2004 Model selection in ecology and evolution. Trends Ecol. Evol. 19, 101–108. (doi:https://doi.org/10.1016/j.tree.2003.10.013)

51. Lundin O, Rundlöf M, Smith HG, Bommarco R. 2012 Towards integrated pest management in red clover seed production. J. Econ. Entomol. 105, 1620–1628.

52. Pywell RF, Heard MS, Woodcock BA, Hinsley S, Ridding L, Nowakowski M, Bullock JM. 2015 Wild-life friendly farming increases crop yield: evidence for ecological intensification. Proc. R. Soc. London. Ser. B, Biol. Sci. 282, 20151740. (doi:10.1098/rspb.2015.1740)

53. Bommarco R, Kleijn D, Potts SG. 2013 Ecological intensification: Harnessing ecosystem services for food security. Trends Ecol. Evol. 28, 230–238. (doi:10.1016/j.tree.2012.10.012)

54. Maes J, Jacobs S. 2015 Nature-Based Solutions for Europe’s Sustainable Development. Conserv. Lett. 10, 121–124. (doi:10.1111/conl.12216)

55. Tamburini G, Lami F, Marini L. 2017 Pollination benefits are maximized at intermediate nutrient levels. Proc. R. Soc. B Biol. Sci. 284.

56. Łukowiak R, Grzebisz W, Sassenrath GF. 2016 New insights into phosphorus management in agriculture — A crop rotation approach. Sci. Total Environ. 542, 1062–1077. (doi:https://doi.org/10.1016/j.scitotenv.2015.09.009)

57. Ijaz M, Mahmood K, Honermeier B. 2015 Interactive Role of Fungicides and Plant Growth Regulator (Trinexapac) on Seed Yield and Oil Quality of Winter Rapeseed. Agronomy 5, 435–446. (doi:10.3390/agronomy5030435)

58. Rose TJ, Rengel Z, Ma Q, Bowden JW. 2008 Post-flowering supply of P, but not K, is required for maximum canola seed yields. Eur. J. Agron. 28, 371–379. (doi:10.1016/j.eja.2007.11.003)

59. Stahl A, Pfeifer M, Frisch M, Wittkop B, Snowdon RJ. 2017 Recent Genetic Gains in Nitrogen Use Efficiency in Oilseed Rape. Front. Plant Sci.. 8, 963.

60. Colnenne C, Meynard JM, Roche R, Reau R. 2002 Effects of nitrogen deficiencies on autumnal growth of oilseed rape. Eur. J. Agron. 17, 11–28. (doi:https://doi.org/10.1016/S1161-0301(01)00140-X)

61. Ozturk O. 2010 Effects of source and rate of nitrogen fertilizer on yield, yield components and quality of winter rapeseed (Brassica napus L.). Chil. J. Agric. Res. 70, 132–141.

62. Cheema MA, Malik MA, Hussain A, Shah SH, Basra SMA. 2001 Effects of time and rate of nitrogen and phosphorus application on the growth and the seed and oil yields of canola (Brassica napus L.). J. Agron. Crop Sci. 186, 103–110. (doi:10.1046/j.1439-037X.2001.00463.x)

63. Rucker RR, Thurman WN, Burgett M. 2012 Honey Bee Pollination Markets and the Internalization of Reciprocal Benefits. Am. J. Agric. Econ. 94, 956–977. (doi:10.1093/ajae/aas031)

64. Gagic V et al. 2017 Combined effects of agrochemicals and ecosystem services on crop yield across Europe. Ecol. Lett. 20, 1427–1436. (doi:10.1111/ele.12850)

65. Glamann J, Hanspach J, Abson DJ, Collier N, Fischer J. 2017 The intersection of food security and biodiversity conservation: a review. Reg. Environ. Chang. 17, 1303–1313. (doi:10.1007/s10113-015-0873-3)

66. Popp J, Pető K, Nagy J. 2013 Pesticide productivity and food security. A review. Agron. Sustain. Dev. 33, 243–255. (doi:10.1007/s13593-012-0105-x)

67. Hiiesaar K, Metspalu L, Lääniste P, Jõgar K, Kuusik A, Jõudu J. 2003 Insect pests on winter oilseed rape studied by different catching methods. Agron. Res. 1, 17–29.

68. Veromann E, Tarang T, Kevväi R, Luik A, Williams I. 2006 Insect pests and their natural enemies on spring oilseed rape in Estonia: impact of cropping systems.

69. Gagic V, Riggi LGA, Ekbom B, Malsher G, Rusch A, Bommarco R. 2016 Interactive effects of pests increase seed yield. Ecol. Evol. 6, 2149–2157. (doi:10.1002/ece3.2003)

70. Lechenet M, Dessaint F, Py G, Makowski D, Munier-Jolain N. 2017 Reducing pesticide use while preserving crop productivity and profitability on arable farms. Nat. Plants 3, 17008.

71. Rader R et al. 2015 Non-bee insects are important contributors to global crop pollination. Proc. Natl. Acad. Sci., 201517092. (doi:10.1073/pnas.1517092112)

72. Rollin O, Bretagnolle V, Fortel L, Guilbaud L, Henry M. 2015 Habitat, spatial and temporal drivers of diversity patterns in a wild bee assemblage. Biodivers. Conserv. 24, 1195–1214. (doi:10.1007/s10531-014-0852-x)

